# Optogenetic interrogation of the role of striatal patches in habit formation and inhibition of striatal dopamine

**DOI:** 10.1101/2020.10.29.359265

**Authors:** J.A. Nadel, S.S. Pawelko, J.R. Scott, R. McLaughlin, M. Fox, N.G. Hollon, C.D. Howard

**Affiliations:** Neuroscience Department, Oberlin College, 173 Lorain St., Oberlin, OH, USA; Molecular Neurobiology Laboratory, The Salk Institute for Biological Studies, 10010 North Torrey Pines Road, La Jolla, CA 92037, USA

## Abstract

Habits are inflexible behaviors that can be maladaptive in diseases including drug addiction. The striatum is integral to habit formation, and interspersed throughout the striatum are patches, or striosomes, which are characterized by unique gene expression relative to the surrounding matrix. Recent work has indicated that patches are necessary for habit formation, but how patches contribute to habits remains partially understood. Here, using optogenetics, we modulated striatal patches in Sepw1-NP67 mice during habit formation. We find that patch activation during operant training impairs habit formation, and conversely, that acute patch stimulation after reward devaluation can drive habitual reward seeking. Patch stimulation invigorates general locomotion but is not inherently rewarding. Finally, we use fast-scan cyclic voltammetry to demonstrate that patch stimulation suppresses dopamine release in dorsal striatum *in vivo*. Overall, this work provides novel insight into the role of the patch compartment in habit formation, and potential interactions with dopamine signaling.

## Introduction

Organisms must optimize action patterns to be successful in their environments. This optimization process can come in two forms: updating of actions can be highly flexible and dependent on outcomes (so-called action-outcome, or goal-oriented behaviors) or, with extended training, action updating can become resistant to change regardless of outcome (stimulus-response or habitual behaviors; Dolan and Dayan, 2013). Habitual, automated behaviors can be highly advantageous, as they allow animals to respond to stimuli without great cognitive effort. However, habits can also present as maladaptive behaviors that persist in spite of negative outcomes. Moreover, dysfunctional habit formation underlies many pathological states, including drug addiction (Robbins and Everitt, 1999).

In animal models, habits have been studied by measuring perseverance of instrumental behaviors following reduction in reward value or by measuring flexibility when action-outcome contingencies are manipulated (Adams and Dickinson, 1981; Dickinson, 1985; Rossi and Yin, 2012). Using these approaches, distinct neural circuits underlying goal-directed and habitual responding have been identified. A well supported model has emerged positing that the dorsomedial striatum encodes goal-directed behaviors, while the dorsolateral striatum encodes habitual behaviors (Yin et al., 2005, 2004; Yin and Knowlton, 2006). Similarly, corticostriatal plasticity in the lateral striatum correlates with habitual responding (O’Hare et al., 2016), and human imaging studies have linked activity in lateral striatum (putamen) with habitual behaviors (Tricomi et al., 2009). However, this model could be somewhat oversimplified, as other studies suggest medial striatum could also contribute to inflexible behaviors (Malvaez et al., 2018; Seiler et al., 2020).

Adding a layer of complexity to the medial-lateral striatal divide is the existence of two neurochemically distinct subcompartments: small, labyrinthine islands called patches or striosomes (comprising 15% of striatal volume), and surrounding ‘matrix’ tissue (85% of striatal volume; Gerfen, 1992; Graybiel and Ragsdale, 1978). In addition to unique cellular markers (Crittenden and Graybiel, 2011), patches are characterized by unique connectivity, providing the predominant anatomical and functional striatal input to midbrain dopaminergic neurons (Evans et al., 2020; Gerfen, 1985). Additionally, habenula-projecting neurons of the entopeduncular nucleus receive preferential input from patches (Stephenson-Jones et al., 2016; Wallace et al., 2017). Striatal patches also have unique input profiles, with preferential inputs from frontal cortex (Eblen and Graybiel, 1995; Gerfen, 1984; but see Smith et al., 2016). Therefore, striatal patches are well-positioned to serve as a limbic-motor interface that could subserve action selection (Shivkumar et al., 2017).

Despite extensive work characterizing the structure and connectivity of striatal patches, their role in behavior regulation is only partially understood. Studies have suggested a role for striatal patches in reward processing (Bloem et al., 2017; White and Hiroi, 1998; Yoshizawa et al., 2018) and cost-benefit decision making (Friedman et al., 2017, 2015). Additionally, several studies now support the notion that patches may encode the transition from flexible to habitual responding. Early studies suggested that psychostimulant-induced stereotypy is linked to activity in patches (Canales and Graybiel, 2000; Murray et al., 2015, 2014) and that lesions of patches disrupt this stereotypy (Murray et al., 2015, 2014). More recently, striatal patches have been shown to be necessary for normal habit formation: specific lesions of patch neurons diminish habitual responding following reward devaluation (Jenrette et al., 2019) or changes in action-outcome contingencies (Nadel et al., 2020).

In the current study, we employed optogenetics in Sepw1-NP67 mice with enriched Cre recombinase expression in striatal patches (Gerfen et al., 2013) to selectively target patches. Patches or patch projections were stimulated at reward retrieval during a variable interval schedule of responding, a task used to induce habitual responding (Gremel and Costa, 2013; Rossi and Yin, 2012). Mice that received stimulation of striatal patches reduced lever pressing and head entry rates to a greater extent than YFP controls following reward devaluation, implying impaired habit formation. Following retraining and subsequent reward devaluation, acute stimulation of patches was sufficient to drive habitual reward seeking behaviors. Contrary to a prior study using non-selective electrical self-stimulation (White and Hiroi, 1998) we did not find optogenetic stimulation of patches to be reinforcing in a place preference task, but stimulation of patches did elevate locomotion in an open field. Finally, to investigate how patch activation modifies circuit function, we employed fast-scan cyclic voltammetry to measure striatal dopamine levels *in vivo* and determined that optogenetic activation of patches suppresses dopamine release driven by electrical stimulation of excitatory inputs. Together, these results suggest striatal patches are a key site underlying habit formation and that activating patches can drive habitual reward seeking, potentially by modulating striatal dopamine levels.

## Results

### Optogenetic manipulation of striatal patches or projections in variable interval training

To investigate the role of patch neurons in habit formation, we utilized Sepw1-NP67 mice, which have enriched Cre recombinase expression in striatal patches (Gerfen et al., 2013; Smith et al., 2016). Crossing these mice with a Cre-dependent GFP reporter line shows enriched GFP+ neurons in μ-opiate receptor dense striatal patches (Crittenden and Graybiel, 2011, Figure 1C), though as previously reported, this line also expressed Cre in “exo-patch” neurons, which display similar gene expression and physiological profiles to patch neurons (Smith et al., 2016). We injected Sepw1-NP67 mice with an AAV encoding either Cre-dependent light-gated cation channel ChR2 or YFP in the dorsal striatum, which resulted in enriched ChR2 expression in striatal patches (Figure 1D). We then implanted fiber optics targeting cell bodies of striatal patch neurons, patch terminals in SNc (Evans et al., 2020), or at patch terminals in entopeduncular nucleus (Stephenson-Jones et al., 2016; Wallace et al., 2017; Figure 1A + B) with the expectation that these two pathways may differentially modulate habitual responding due to potentially opposing effects on dopamine neurons. However, no implantation site-dependent differences were observed in performance during training, habit probes, open field, or place preference tasks (p > 0.05), therefore fiber optic placement groups were collapsed into a general “ChR2” group for comparison with YFP controls (individual group data is shown in supporting figures matching main figure numbers; Supporting Figures 2–5).

**Figure 1.**
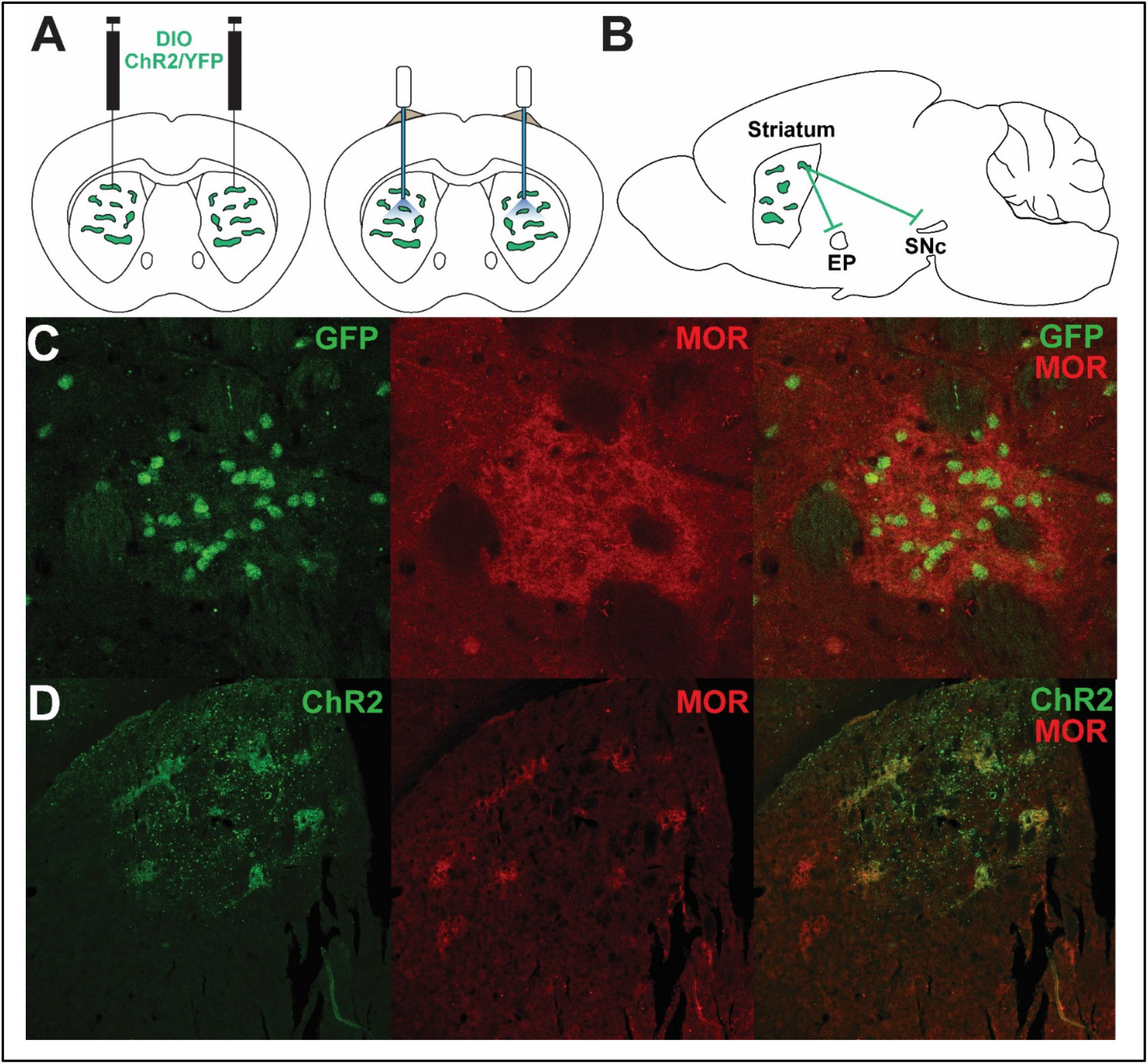
Experimental Design and Characterization of Sepw1–Cre Mice. A. Experimental design. Bilateral AAV driving expression of channelrhodopsin-2 (ChR2) or YFP was injected into dorsal striatum (left). Fiber optics were affixed just dorsal to injection site (right). B. Patches in striatum, or patch projections to entopeduncular nucleus (EP) or substantia nigra pars reticulata were targeted with fiber optic implants. C. Coronal section showing Cre dependent expression of GFP and μ-opioid receptor expression demonstrating enriched Cre expression in a striatal patch. D. Coronal section showing AAV5-driven expression of ChR2-eYFP overlaid with μ-opioid receptor expression.

Three weeks after surgery, mice were food restricted and trained to depress a lever on a continuous reinforcement schedule (CRF), a variable interval averaging 30 sec (VI30), then a variable interval averaging 60 sec (VI60) schedule of reinforcement (see Figure 2A for behavioral schedule), which induces habitual behavior in mice (Rossi and Yin, 2012). Beginning in VI60, mice received laser stimulation through fiber optics at reward retrieval (first headentry following reward delivery; 3 sec, 5 Hz, 5 mW stimulation). Both ChR2 and YFP mice increased press rates across CRF and VI training (two-way repeated measures ANOVA, significant effect of time, F_(14, 392)_ = 20.04, p < 0.0001). However, ChR2-stimulated mice had a tendency to press at a slightly higher rate across training (trending effect of group, F_(1, 28)_ = 3.841, p = 0.060; no significant interaction, p > 0.05; Figure 2B). This phenomenon was not attributable to differences during VI30 training (t_28_ = 0.7205, p = 0.477; Figure 2C), but rather a tendency for ChR2 mice to increase pressing following the onset of stimulated trials in VI60 (t_28_ = 2.013, p = 0.054; Figure 2D). We previously found that caspase-driven lesions of striatal patches increased response variability across days (Nadel et al., 2020), and similarly, optogenetic activation of patches during VI60 training had a tendency to reduce press rate consistency as assessed by day-to-day autocorrelation (lag 1 day; unpaired t-test, t_28_ = 1.691, p = 0.102; Figure 2E).

**Figure 2.**
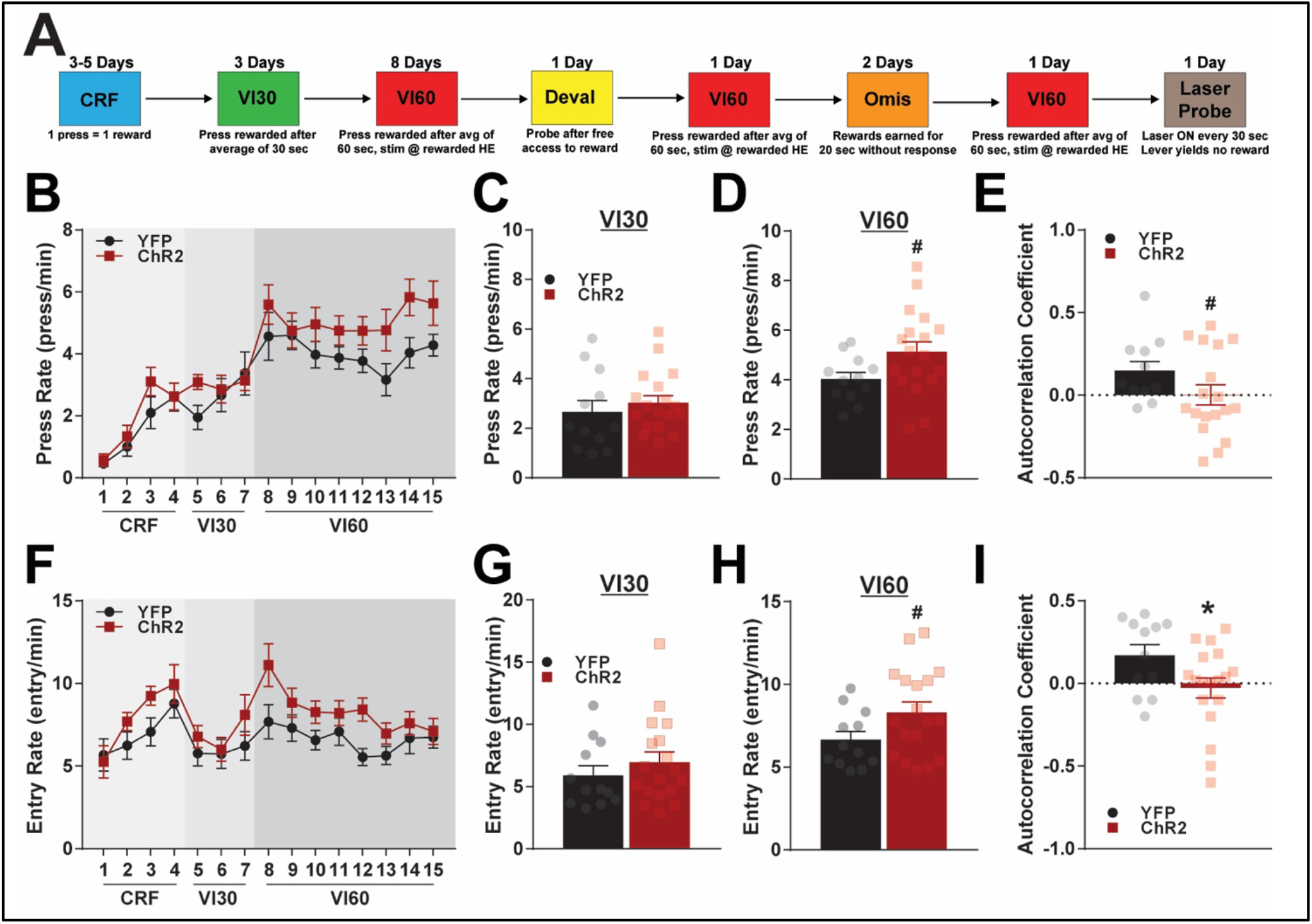
Optogenetic patch stimulation during variable interval training. A. Timeline of behavioral schedule. Mice were trained on a continuous reinforcement schedule (CRF) before beginning variable interval 30 training (VI30). Following this, mice began a variable interval 60 schedule (VI60), and optogenetic stimulation of patches or patch projections occurred during reward retrieval. Mice then experienced reward devaluation before being retrained on VI60. Mice then experienced and two days of omission (Omis). Finally, following retraining on VI60, mice experienced a “Laser Probe” trial. See Methods for details of each behavioral schedule. B. Press rates for ChR2 and YFP control mice across training. C-D. Average response rates across VI30 (C) or VI60 (D). E. Autocorrelation coefficients for press rates across VI60 training. F. Head entry rates to the food magazine across training for ChR2 and YFP mice. G-H. Average head entry rates across VI30 (G) and VI60 (H). I. Autocorrelation coefficients for entry rates across VI60 training. *\*P* < 0.05; #*P* < 0.1; error bars, SEM.

In addition to lever pressing, head entry to the food magazine has been used to assess flexibility (DePoy et al., 2016; Morrison et al., 2015; Sieburg et al., 2019). ChR2 mice and YFP controls both altered their entry rates across training (two-way repeated measures ANOVA, significant effect of time, F_(14, 392)_ = 5.148, p <0.0001), though ChR2 mice tended to have higher entry rates (trending effect of group, F_(1, 28)_ = 3.204, p = 0.0843; no significant interaction, p > 0.05; Figure 2F). This was also not attributable to VI30 (t_28_ = 0.8953, p = 0.3782 Figure 2G), but a trending increase in entry rate was noted during stimulated VI60 trials (t_28_ = 1.922, p = 0.0649; Figure 2H). Additionally, activating patches across VI60 training resulted in even clearer disruption of day-to-day consistency in head entries as assessed by autocorrelation (lag 1 day; unpaired t-test, t_28_ = 2.145, p = 0.0407; Figure 2I). Taken together, this data suggests that ChR2 injections do not impair learning in CRF or VI30, but that stimulation of patches slightly invigorates responding in VI60 when stimulation is paired to rewarded head entries. Further, this data suggests that modulation of patches impairs day-to-day response consistency.

### Characterizing patches stimulation during learning in devaluation and omission probes

Habits are operationally defined as behaviors resistant to outcome devaluation. Therefore, after the completion of eight VI60 training days, mice received free access to sucrose for an hour before being returned to behavioral chambers for a 5 min devaluation probe trial. Visual confirmation was made to ensure each mouse drank sucrose during free access, and the weight of sucrose consumed was not significantly different between implantation sites (Fig, S3A; p > 0.05) or between ChR2 and YFP mice (Figure S3B; p > 0.05) in a subgroup of mice. During probe trials, lever presses and head entries were recorded, but no rewards were delivered. ChR2 and YFP mice did not significantly differ in raw press rate during devaluation (unpaired t-test, t_28_ = 1.382, p = 0.178; Figure 3A). However, due to the increased variability in ChR2 mice and slightly different press rates between groups, we normalized the devaluation press rate to mean press rates across VI60 for each mouse. ChR2 mice pressed significantly less in the devaluation probe when normalized to baseline responding, indicating weaker habit formation (unpaired t-test, t_28_ = 2.261, p = 0.0317; Figure 3B). Similarly, ChR2 mice entered the reward port less frequently than YFP controls during devaluation probes, both in raw (t_27_ = 3.398, p = 0.0021; Figure 3C) and normalized entry rate (t_27_ = 3.845, p = 0.0007; Figure 3D).

**Figure 3.**
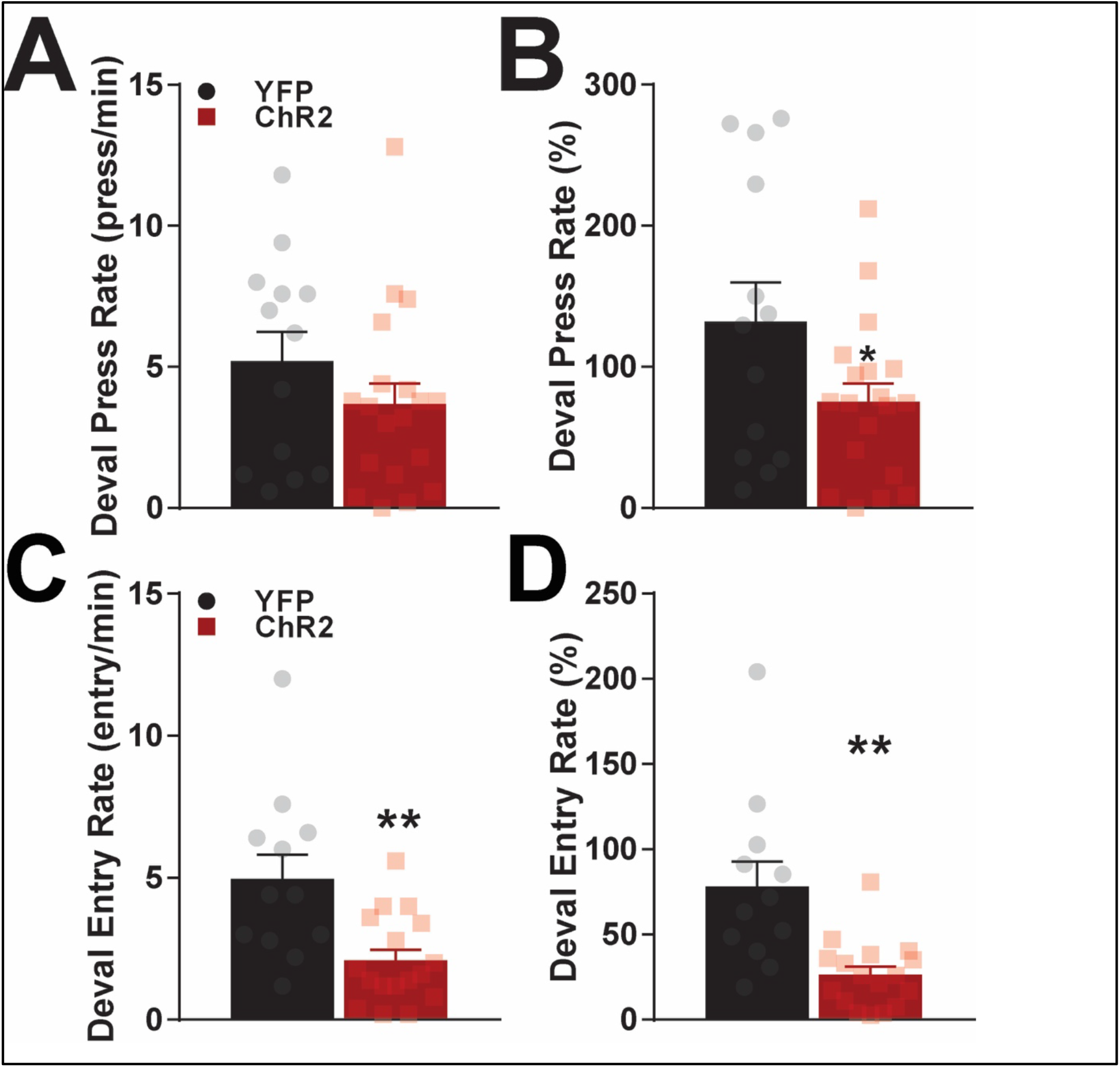
Effects of optogenetic patch manipulation in learning following devaluation. A. Average press rates during devaluation probe trials (see Methods). B. Average press rates in devaluation probes normalized to baseline responding during VI60 training. C. Average head entry rates during devaluation probe trials. D. Average head entry rates in devaluation probes normalized to baseline entry rate during VI60 training. *\*P* < 0.05; error bars, SEM.

One day after devaluation probes, mice were retrained on a VI60 schedule with optogenetic stimulation of patches to reinstate robust pressing before beginning two days of omission probes. In omission, mice were required to abstain from pressing for 20s in order to receive a reward. This probe has been used as an alternative means to assess habit by measuring flexibility in forming new action-outcome contingencies (Yin et al., 2005, 2004), and we previously reported that mice with patch lesions have reduced pressing in omission trials. However, we did not observe any effect of ChR2 stimulation on raw or normalized press or entry rates relative to controls (Supporting Figure 1A-F, all p > 0.05). As expected, YFP mice had a strong correlation between press rate on day 1 of omission and press rate on the reinstatement day between devaluation and omission probes (Pearson’s correlation, R^2^ = 0.551, p = 0.0057; Supporting Figure 1G). However, ChR2 mice did not display any correlation between press rates during these days (R^2^ = 0.0095, p = 0.6993; Supporting Figure 1H). This finding could further suggest impaired day-to-day consistency in responding in patch stimulated mice.

### Determining acute effects of optogenetic stimulation of patches following devaluation

If patches encode habits, we reasoned that acute optogenetic stimulation of patches may drive habitual responding even following reward devaluation. Mice therefore underwent another day of VI60 retraining with optogenetic stimulation to reinstate robust pressing. Following this, mice began a novel probe trial, which began with one hour of free access to sucrose to devalue rewards. Mice were then placed in the operant chamber with the lever extended, though no rewards were delivered. Laser stimulation was delivered on a variable 60 sec interval, which was not contingent on responding, and trials lasted 30 min (referred to as a “laser probe” trials). Press rates for YFP and ChR2 are shown relative to laser onset (blue) in Figure 4A. Laser stimulation did not alter pressing behaviors in YFP mice (paired t-test, t_8_ = 0.0622, p = 0.9519; Figure 4B). To our surprise, laser stimulation in ChR2 mice was immediately followed by a near-significant decrease in lever pressing (t_10_ = 2.193, df = 10, p = 0.0531; Figure 4C). To investigate what happens during this decrease in pressing, we repeated this analysis focusing on head entries before and after stimulation (Figure 4D). Again, stimulation in YFP controls did not alter entry rate (t_8_ = 0.4415, p = 0.6705), however, ChR2 immediately increased entry rate following stimulation (t_10_ = 3.049, p = 0.0123). These data suggest that patch stimulation drives habitual reward seeking following reward devaluation.

**Figure 4.**
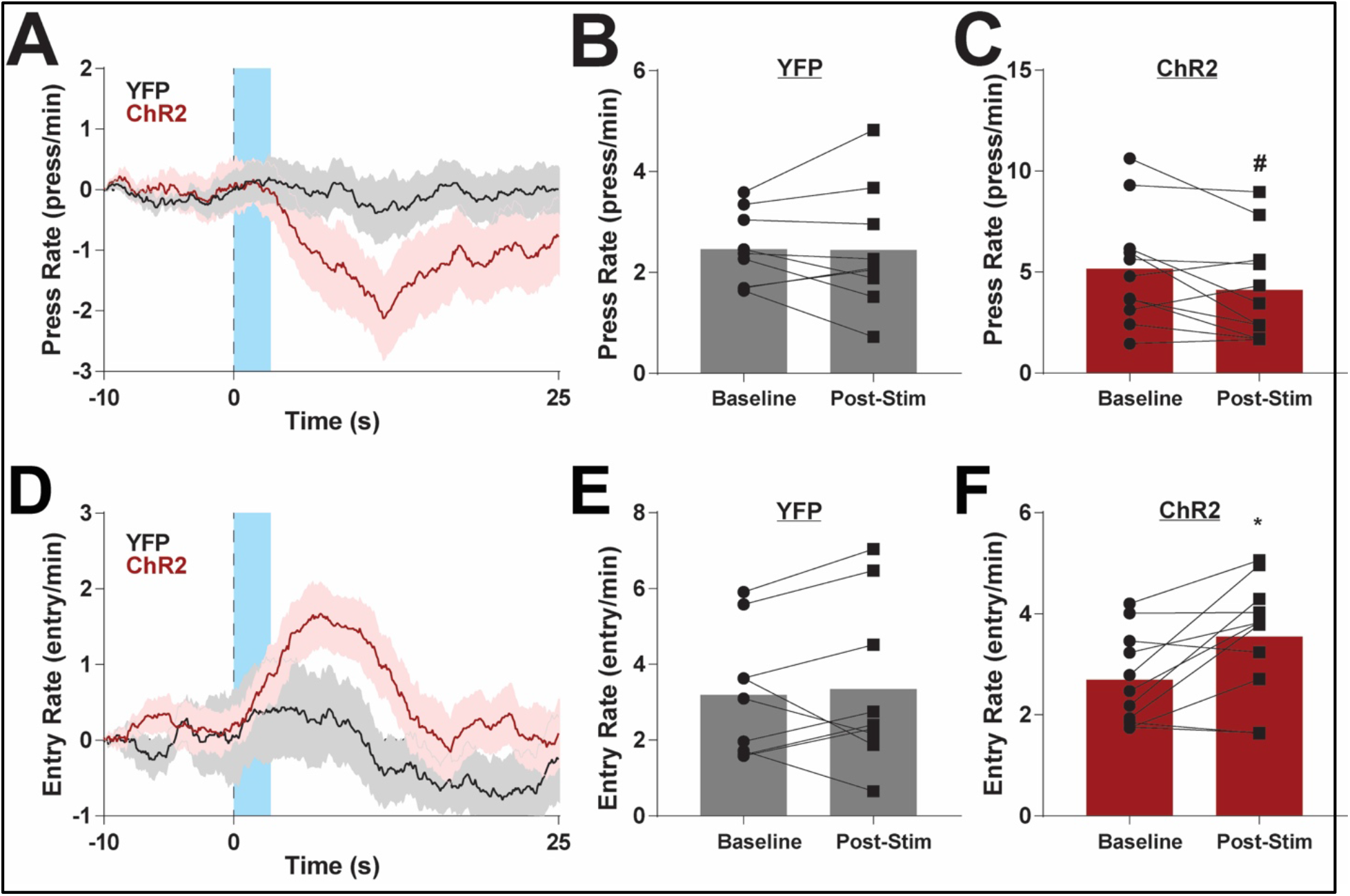
Effects of acute optogenetic patch manipulation in laser probe trials. A. Average baseline normalized press rates before and after laser onset. Laser stimulation (5Hz, 3 sec) is indicated by blue. B-C. Press rates before and after laser onset for YFP (B) or ChR2 (C) mice. D. Average baseline normalized head entry rates before and after laser onset. E-F. Head entry rates before and after laser onset for YFP (E) or ChR2 (F) mice. \**P* < 0.05; #*P* < 0.1; error bars, SEM.

### Determining patch contributions to locomotion and place preference

Patch stimulation could modify responding by being inherently rewarding (White and Hiroi, 1998). To explore this possibility, we next investigated the effects of patch stimulation on reinforcement in a place preference task. To test the effects of optogenetic stimulation on reinforcement, mice began a two-day real-time place preference task in a 2-chamber apparatus. Entry to a randomly selected side resulted in laser stimulation (5 sec ON and 5 sec OFF, cycled), which ended upon entrance to the opposite chamber. The stimulation side was counterbalanced across 2 days and preference was averaged between days (see Methods). Patch activation did not drive differences in time spent on the stimulation side relative to YFP controls (unpaired t-test, t_27_ = 1.143, p = 0.2631; Figure 5A). These results suggest that optogenetic stimulation of patches is not inherently reinforcing in this place preference task.

**Figure 5.**
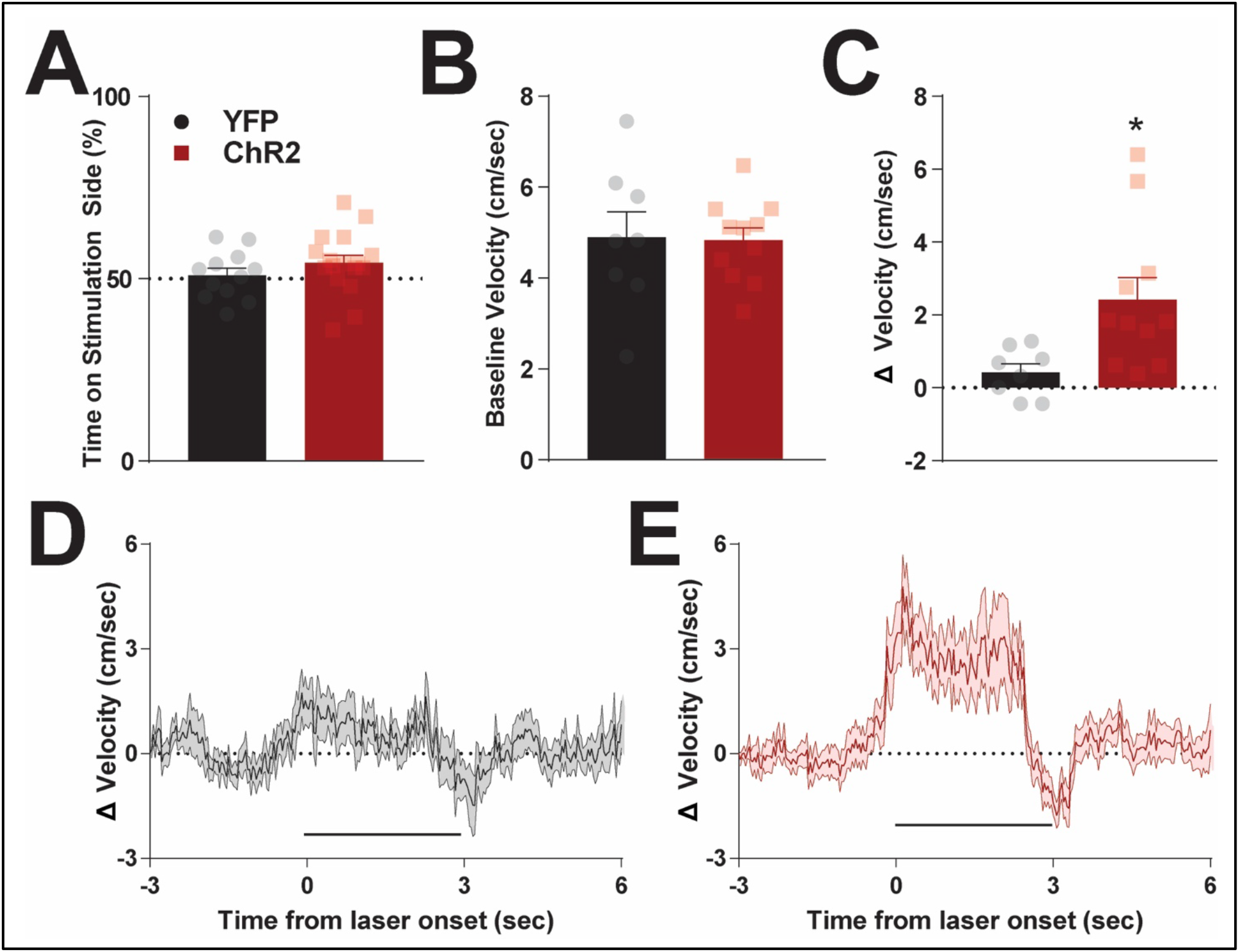
Effects of optogenetic stimulation of patches on reinforcement and locomotion. A. Time spent on stimulated side of a two-chamber place preference apparatus. Time is averaged between two days, and stimulated side was counter balanced between days (see Methods). B. Average velocity in the open field for YFP and ChR2 mice. C. Change in velocity following laser onset in open field. D-E. Average baseline normalized velocity before and after laser onset (5 Hz, 3 sec; denoted by thick black line) for YFP (D) and ChR2 (E) mice. \**P* < 0.05; error bars, SEM.

Due to a predominant D1 dopamine receptor makeup (Smith et al., 2016), patch stimulation may alter behavior by invigorating movement (Kravitz et al., 2010). To explore this possibility, mice were placed in a custom open field chamber to assess the effects of patch stimulation on locomotion, and laser stimulation occurred every 60 sec while the location of the mouse was tracked (see Methods). We first assessed if intermittent patch stimulation elevated average locomotion throughout the task, and found no difference in movement speed between ChR2 and YFP mice (unpaired t-test, t_17_ = 0.1127, p = 0.9116; Figure 5B). However, onset of laser stimulation significantly increased locomotion in ChR2 mice relative to controls (unpaired t-test, t_17_ = 2.708, p = 0.0149; Figure 5C). On a finer timescale, YFP controls showed modest responses to laser stimulation (Figure 5D). On the other hand, ChR2 mice demonstrated robust initial increases following laser onset, which plateaued until cessation of stimulation, followed by a short reduction in movement (Figure 5E). Together, these results show that patch activation can acutely drive locomotion without being intrinsically reinforcing.

### Characterization of patch and dopamine interactions *in vivo*

As striatal patches are a unique striatal output population projecting to SNc (Crittenden et al., 2016; Davis et al., 2018; Evans et al., 2020), optogenetic stimulation of patches may alter responding by modulating dopamine release. We aimed to investigate this possibility by eliciting phasic increases in striatal dopamine with electrical stimulation of the pedunculopontine tegmental nucleus (PPTg; Forster and Blaha, 2003; Zweifel et al., 2009) with and without simultaneous optogenetic activation of patch projections. We first injected a group of Sepw1-NP67 mice with AAV encoding Cre-dependent ChR2. Three weeks later, mice were anesthetized and a glass-sealed carbon-fiber microelectrode capable of detecting real-time changes in dopamine with fast-scan cyclic voltammetry was lowered into the dorsal striatum. Following this, a bipolar stimulating electrode was placed in the PPTg. Once robust dopamine was detected, a fiber optic was lowered into the SNc targeting patch terminals (see Figure 6A for experimental design).

**Figure 6.**
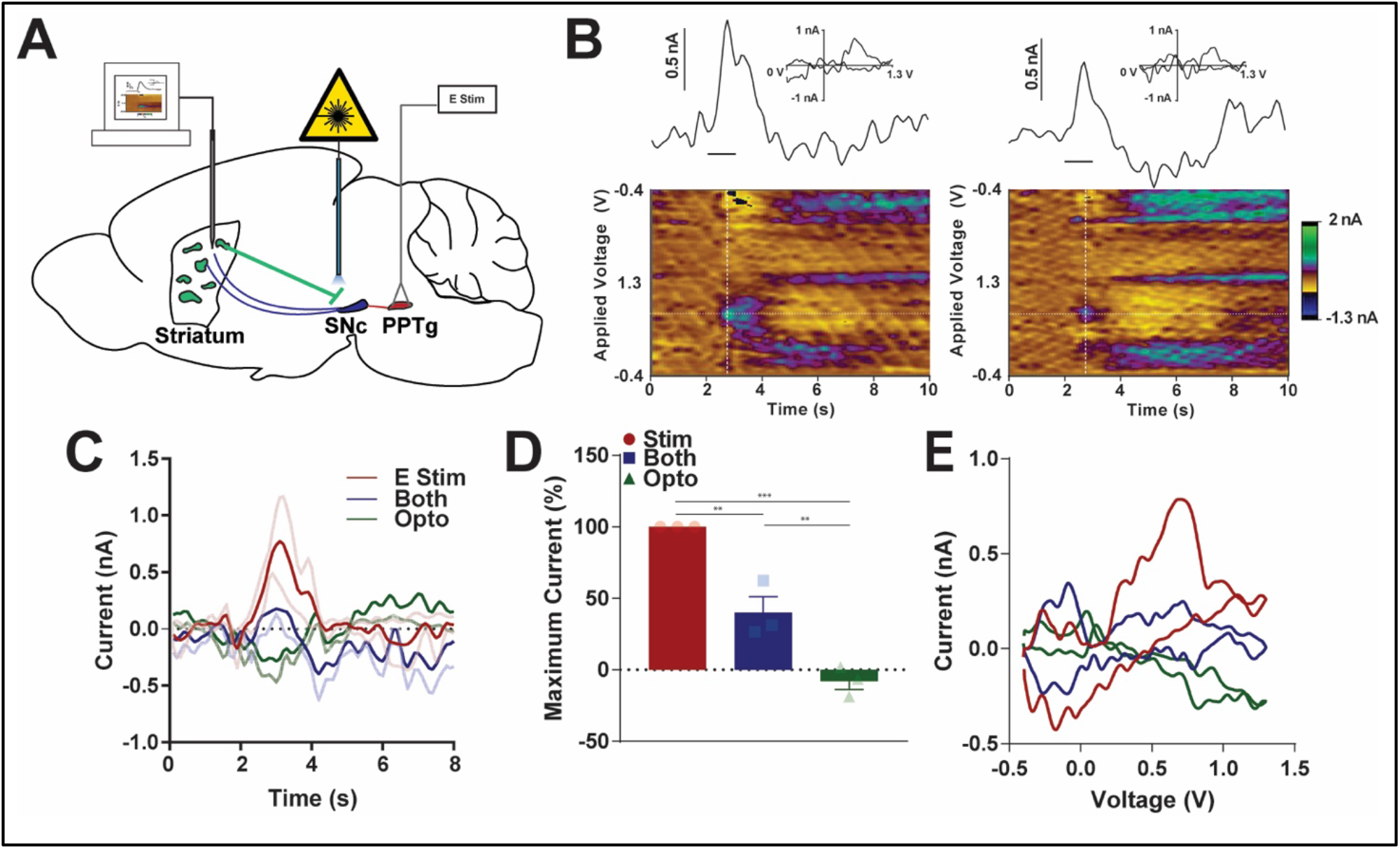
Characterizing optogenetic patch stimulation on striatal dopamine release. A. Experimental design. Sepw1 NP67 mice were injected with AAV5 driving Cre-dependent expression of ChR2-eYFP. Fast-scan cyclic voltammetry was used to monitor real-time changes in dopamine levels in striatum. An electrical stimulating electrode was placed in pedunculopontine nucleus (PPTg) to drive increases in dopamine in dorsal striatum. A fiber optic was placed above patch terminals in substantia nigra pars compacta (SNc). Electrical or optogenetic or both stimulation types were randomly selected and delivered a minimum of three times each (see Methods). B. (left) Representative recording of electrical stimulation of PPTg. Here, a line shows recorded current relative to stimulation delivery (straight line below current trace) above a pseudo-color plot. The color plot shows current collected (in color) at each waveform scan (y-axis) and across time (x-axis). INSET: a “cyclic voltammogram” collected at the vertical white dotted line on the pseudo-color plot suggesting that dopamine is the predominate analyte being monitored. (right) Same as left, but optogenetic stimulation of patch terminals occurs simultaneously with PPTg electrical stimulation. C. Average responses across replicates for each of the three stimulation types. PPTg stimulation only denoted as E Stim, optogenetic stimulation of patch terminals as Opto. D. Maximum recorded current during stimulation normalized to electrical PPTg stimulation. E. Average cyclic voltammogram across replicates for each stimulation type. \**P* < 0.05; \*\**P* < 0.01; \*\*\**P* < 0.001; error bars, SEM.

Stimulation of PPTg resulted in increases in striatal dopamine that mimicked naturally occurring phasic increases in dopamine (Howard et al., 2013; Robinson et al., 2002; Figure 6B, left). When PPTg stimulation occurred simultaneously with optogenetic patch activation, phasic dopamine responses were present, but reduced in amplitude (Figure 6B, right). On average, PPTg stimulation alone resulted in larger responses relative to simultaneous electrical and optogenetic stimulation. On the other hand, optogenetic patch activation alone caused a small decrease in detected current (Figure 6C; Supporting Figure 6A). When recordings were normalized to average PPTg stimulation recording amplitude to account for baseline differences in release between subjects, PPTg stimulation drove a larger dopamine response than simultaneous PPTg and patch activation, which was significantly higher than optogenetic activation alone (One-way ANOVA, F_(2,6)_ = 53.97, p < 0.0001;*post hoc* Tukey multiple comparison test all p < 0.01; Figure 6D).

Inspection of average cyclic voltammograms further suggests the analyte detected following PPTg stimulation is dopamine, and the peak oxidation is reduced following simultaneous optogenetic and electrical stimulation (Figure 6E; average current and cyclic voltammograms for each experimental replicate shown in Supporting Figure 6B). These results suggest that patch projections to dopamine neurons are capable of suppressing dopamine release in the dorsal striatum, which may contribute to the effects noted in the behavioral tasks.

## Discussion

The striatum is a key locus in the transition from flexible to habitual strategies, but less is known about how particular striatal subcircuits contribute to this phenomenon. Previous studies have implicated striatal patches, or striosomes, in this process (Canales and Graybiel, 2000; Murray et al., 2015, 2014), and more recent work has demonstrated that intact patches are necessary for normal habit formation (Jenrette et al., 2019; Nadel et al., 2020). The current work further addresses the role of patches in habit through use of optogenetics to modulate patch neuron activity in a temporally-precise manner during habit formation. Patch stimulation enhanced behavioral variability and invigorated responding across training. Additionally, optogenetic stimulation of patch activity during learning suppressed the rate of lever pressing and head entry to the reward port following reward devaluation, suggesting impaired habit formation. Next, we developed a novel probe trial aimed at determining how mice respond following stimulation of patches following reward devaluation. In these so-called ‘laser probe’ trials, stimulation of patches acutely suppressed lever pressing and augmented head entry to the food port, suggesting acute patch activation can drive habitual reward seeking. These effects were likely not attributable to directly reinforcing effects of patch stimulation as assessed in a place preference task, though stimulation of patches was sufficient to augment locomotion in an open field. Finally, we demonstrated that optogenetic stimulation of patch terminals is sufficient to suppress dopamine release driven by electrical stimulation of excitatory inputs to dopamine neurons. Together, these results suggest that patches are a key site of habit formation and that patch activation can modify habitual responding, potentially through regulation of striatal dopamine levels.

### Patches as a locus of habitual behavior

The finding that patch manipulation impairs habitual responding is supported by previous studies using Cre-dependent caspase lesions (Nadel et al., 2020) or a conjugated cytotoxin that selectively ablates μ-opioid receptor expressing neurons (Jenrette et al., 2019), both finding impairments in habitual responding. These patch manipulations, and the optogenetic approach utilized here, target the central striatum, likely affecting patches in both medial and lateral striatum. A now well-supported model proposes that the medial striatum encodes goal-oriented behavior, while the lateral striatum encodes habitual responding (Yin and Knowlton, 2006), a distinction which may also apply to dopamine neurons (Faure et al., 2005; Lerner et al., 2015). Patches span the medial/lateral spectrum of the striatum, and studies manipulating patches specifically within medial or lateral striatum should be pursued to determine if there is a functional divide within medial and lateral patches.

One puzzling aspect of the current work is how activation of patches through the use of optogenetics could impair habit formation, when lesioning patches has a similar effect (Jenrette et al., 2019; Nadel et al., 2020). We propose two ideas to explain this paradoxical finding. First, optogenetic stimulation may enhance ongoing patch activity, but could impair the timing of spiking relative to afferent activation during learning, thus disrupting plasticity during the transition to habitual responding. Striatal neurons have been shown to be highly sensitive to spike-timing of corticostriatal inputs, and modifying afferent and spiny projection neuron spike timing by milliseconds can reverse the valence of plasticity in these cells (Fino and Venance, 2010). We chose to deliver optogenetic stimulation during reward retrieval based on previous studies showing activity increases in patches to rewards or cues predicting rewards (Bloem et al., 2017; Yoshizawa et al., 2018). However, direct electrophysiological or optical recordings of patches would be required to determine patch activity in this variable interval schedule. Alternatively, optogenetic activation of patches may drive rebound inhibition, which may lead to impairments in habit formation by suppressing patch activity following cessation of laser firing. Rebound excitation and inhibition have been well characterized during inhibition or excitation of neuronal circuits with optogenetics (Häusser, 2014). Indeed, we note potential behavioral evidence of this phenomenon in the current work: patch activation in open field drives robust increase in locomotion, followed by a brief inhibition of movement where locomotor behaviors fall below baseline (Figure 5E). Future physiological studies of patch function should explore how patch activity is modified during habit learning.

### Patch manipulation decreases responding in devaluation probe trials

Reduced responding during devaluation probes could be partially explained by an elevated response rate during stimulated VI60 trials. That is, if laser stimulation invigorates responding, normalizing responding to an elevated baseline may drive this effect. While this possibility cannot be completely resolved, a lower number of total head entries, which are not normalized to baseline (Figure 3C), argues against this being the only factor contributing to reduced responding following devaluation. An additional limitation of the current study is the lack of a matrix specific manipulation to determine specificity of this effect is to patches. Indeed, matrix lesions have been shown to impair fine motor coordination (Lopez-Huerta et al., 2016), but the role of the matrix has not been investigated in habit formation. However, this work suggests that specific manipulation of patch neurons is sufficient to alter habit formation, adding to a growing body of literature indicating patches are a key site in the transition to habitual responding (Canales and Graybiel, 2000; Jenrette et al., 2019; Murray et al., 2015, 2014; Nadel et al., 2020).

Importantly, the current work lacks a “non-devalued”, or “valuation” probe trial, which controls for general satiety following free access to rewards (Balleine and Dickinson, 1998; Colwill and Rescorla, 1985). Valuation probes often utilize free access to a reward that is different from rewards earned during in training (eg. maltodextrin solution, homecage chow; Nelson and Killcross, 2006; Shillinglaw et al., 2014; Yu et al., 2009) and several studies have described the degree of habit as a ratio of responding in devaluation vs. valuation probe trials (Gremel and Costa, 2013; O’Hare et al., 2017, 2016). Valuation probes were omitted in the current study due to our previous finding that Sepw1-NP67 mice rapidly suppress responding across two days of probe trials regardless of reward type, which obscured results of devaluation (Nadel et al., 2020). When sucrose was weighed before and after free access in a subgroup of mice, we found no difference in the weight of sucrose consumed (Figure S3A+B) suggesting similar satiety across animals. Nevertheless, lack of a control to ensure reward-specific devaluation is a potential confound of the current study.

Stimulation of patches across training reduced head entry to the food magazine in devaluation trials. On the other hand, acute activation of patches following reward devaluation was sufficient to drive head entry to the food magazine. This unexpected finding could suggest that patch activation directly drives habitual reward seeking. Previous studies have utilized head entry as a metric for habitual responding and reward devaluation has been shown to reduce head entry in goal-directed animals (DePoy et al., 2016; Morrison et al., 2015; Rode et al., 2020). Discrete head entry and lever pressing events could be ‘chunked’ into larger learned action sequences that are reinforced across habit formation (Dezfouli and Balleine, 2013). Striatal patches may serve as a neural substrate of this hierarchical reinforcement, as both press and entry rate are reduced in devaluation probes following patch manipulation. Because striatum is necessary for expression of learned action sequences (Berridge and Whishaw, 1992; Yin, 2010), and as striatal activity encodes action chunking (Jin et al., 2014; Jin and Costa, 2010) with differential contribution of direct and indirect pathways (Geddes et al., 2018), future studies should explore the contribution of patch/matrix subcircuits on sequence learning.

### Patches and behavioral variability

During training, optogenetic stimulation of patches resulted in lower autocorrelation coefficients (Figure 2E+I), suggesting impaired day-to-day consistency in responding. Further, correlation of responding during retraining and omission probes were disrupted in ChR2 mice (Supporting Figure 1G-H) which may reflect enhanced behavioral variability across days. These findings are supported by our previous study, which found that Cre-dependent lesions in Sepw1-NP67 mice similarly disrupted autocorrelations and increased behavioral variability (Nadel et al., 2020). These studies together suggest that patches may support habit formation by facilitating crystallization of action patterns. Indeed, generalized lesions of the dorsal striatum tend to increase behavioral variability in foraging tasks (Charnov, 1976; Compton, 2004). In support of this notion, mice in “laser probe” trials show a correlation between response rates in retraining and following devaluation (Supporting Figure 4 K-L). It is possible that patch neurons may play a general role in reducing behavioral variability across learning, though more studies are required to directly test this idea.

### Patches and behavioral invigoration

Additionally, in training, we found that optogenetic stimulation of patches tended to enhance ongoing behaviors. ChR2 mice showed slightly increased press and entry rates during VI60 (Figure 2D+H), increased locomotion in open field (Figure 5C-E), and drove increased entry into the food port during laser probes (Figure 4D+F). This may be due to an enriched population of direct-pathway, D1 dopamine receptor-expressing neurons in patches (Miyamoto et al., 2018; Smith et al., 2016). Indeed, optogenetic stimulation of D1 populations enhances movement (Kravitz et al., 2010) through inhibition of basal ganglia output nuclei (Freeze et al., 2013).

However, a recent study suggests that striosomes can be further subdivided into functionally distinct populations, both of which predominantly express D1 receptors. Contrary to the current work, optogenetic stimulation of Teashirt family zinc finger 1 (Tshz1) expressing neurons in striosomes drives aversion and suppression of movement. On the other hand, optogenetic activation of prodynorphin expressing neurons, which are also enriched in patches drives reinforcement and activation of movement (Xiao et al., 2020). Based on this, it is possible that our Sepw1 NP67 line overlaps more closely with prodynorphin-Cre mice, which is supported by previous work (Smith et al., 2016). Future studies will undoubtedly move toward further subdividing diverse neuron populations in patches and matrix to determine their role in behavioral regulation.

### Patch-dopamine interactions

This work provides new insight into the relationship between striatal patches and dopamine release, demonstrating that optogenetic stimulation of patch projections suppresses dopamine release in the dorsal striatum (Figure 6). Previous studies have supported this notion demonstrating anatomical (Crittenden et al., 2016; Fujiyama et al., 2011; Gerfen, 1985; Watabe-Uchida et al., 2012) and functional connectivity (Evans et al., 2020; McGregor et al., 2019). Patches could therefore regulate habitual behavior by sculpting dopamine release across learning. Indeed, dopamine responses transition from ventromedial to dorsolateral striatum as behaviors become well learned (Willuhn et al., 2012), and patches could be involved in gating of dopamine release early in learning. Moreover, patches could provide feed-forward inhibition to dopamine neurons, which may facilitate the activity shift from reward to cue during Pavlovian conditioning (Schultz, 1998), and which could drive negative dopamine responses during reward omission (Watabe-Uchida et al., 2017). Very recent work suggests that patch-dopamine interactions are also reciprocal, as dopamine differentially modulates patch neuron activity relative to matrix neurons (Prager and Plotkin, 2018). Based on this proposed role of patches regulating dopamine release and habitual behaviors, it will be of great interest to explore how patches contribute to pathological compulsive states including drug addiction and Obsessive Compulsive Disorder.

## Materials and methods

### Key Resources Table

|  |  |  |  |  |
| --- | --- | --- | --- | --- |
| strain, strain background (*Mus musculus*) | Sepw1-NP7-Cre | Charles Gerfen (National Institutes of Health) and Nathaniel Heintz (Rockefeller University) (Gerfen *et al.* 2013) | RRID:SCR_011431 |  |
| strain, strain background (*Adeno-associated virus*) | AAV-EF1a-DIO-hChR2(H134R)-EYFP-WPRE-pA | UNC Viral Vector Core | RRID:SCR_002448 | 4x10<sup>12</sup> particles per ml |
| strain, strain background (*Adeno-associated virus*) | AAV-EF1a-DIO-EYFP | UNC Viral Vector Core | RRID:SCR_002448 | 3.5x10<sup>12</sup> particles per ml |
| antibody | anti-MOR antibody | Immunostar (24216) | RRID:AB_572251 | 1::1000 |
| antibody | 2° antibody | Jackson Laboratory (711-165-152) | RRID:AB_2307443 | 1::250 |

### Animals

All experiments were in accordance with protocols approved by the Oberlin College Institutional Animal Care and Use Committee. Mice were maintained on a 12 hr/12 hr light/dark cycle and unless otherwise noted, were provided *ad libitum* access to water and food. Experiments were carried out during the light cycle using 41 heterozygous Sepw1-Cre^+/-^ mice ranging from 2 to 6 months of age, which were generously provided by Charles Gerfen (National Institutes of Health) and Nathanial Heintz (Rockefeller University). These mice preferentially express Cre-recombinase in striatal patches (Gerfen et al., 2013; Smith et al., 2016, Figure 1C+D).

### Reagents

Isoflurane anesthesia was obtained from Patterson Veterinary (Greeley, CO, USA). Sterile and filtered phosphate buffered saline (PBS, 1X) was obtained from GE Life Sciences (Pittsburgh, PA, USA). Unless otherwise noted, all other reagents were obtained through VWR (Radnor, PA, USA).

### Stereotaxic Surgery and Viral Injections

Sepw1-NP67 mice were anaesthetized with isoflurane (4% at 2 L/sec O2 for induction, 0.5-1.5% at 0.5 L/sec O2 afterward) and then placed in a stereotactic frame (David Kopf Instruments, Tajunga, CA, USA). The scalp was sterilized with povidone iodine and an incision was made in the scalp. For optogenetic experiments, the skull was scored with Optibond (Patterson Dental). Holes were then drilled bilaterally above the dorsal striatum (+0.9 AP, 1.8 ML, −2.5 DV) and 500 nL of an AAV encoding channelrhodopsin (ChR2) (AAV-EF1-DIO-hChR2(H134R)-EYFP-WPRE-pA, UNC Viral Vector Core) was injected. Control mice were injected with an AAV encoding YFP (AAV-EF1a-DIO-EYFP, UNC Viral Vector Core). For all injections, a 5 μL syringe needle (Hamilton) was lowered to the DV coordinate over 2 minutes and held in place for 1 min before the start of injection. The injection speed was 100 nL/min, and the needle was left undisturbed in the brain for 5 minutes after the completion of virus delivery, after which the needle was removed over the course of 5 minutes. Fiber optics were then inserted bilaterally targeting one of three sites: cell bodies of patch neurons in the striatum (+0.9 AP, 1.8 ML, −2.3 DV), patch terminals at dopamine neurons of the substantia nigra pars compacta (−3.2 AP, 1.5 ML, −3.6 DV), or over patch terminals in the entopeduncular nucleus EP (−1.1 AP, 2.1 ML, −4.0 DV; Figure 1C+D), and secured to the skull with dental cement (Patterson Dental). Control mice expressing YFP had fiber optics implanted targeting one of these three sites selected randomly. Carprofen (5 mg/kg, subcutaneous) was used for postoperative analgesia. A subset of mice were injected with AAV encoding ChR2 but did not receive fiber optic implants. These mice instead received sterile sutures to close the incision site (see Fast-Scan Cyclic Voltammetry below). All mice were given 3-4 weeks to allow for viral expression and to recover before behavioral training started.

### Variable Interval Training

Mice were trained on a variable interval schedule to induce habitual responding (Rossi and Yin, 2012, see Figure 2A for schematic of entire behavioral training protocol). Throughout training, mice were food deprived and kept at ~85% initial weight by daily feeding of 1.5-2.5g of homecage chow daily after training. All instrumental learning experiments were performed in standard operant chambers (Med Associates). Each chamber had a retractable lever on either side of a reward bowl, which was linked to a sucrose-filled syringe that delivered liquid reward (10% sucrose solution, 20 μl) and a house light on the opposite side of the chamber. Briefly, mice first underwent four days of continuous reinforcement (CRF, one lever press yields one reward) to establish the association between lever press and reward. At the start of the session, the house light was illuminated, and one lever was inserted into the chamber. After 60 min or 50 rewards, the light was shut off, the lever was retracted, and the session ended. On the final day of CRF training, mice were briefly anesthetized with isoflourane (4%, 2 l/min O2) and were connected to fiber optic leads to habituate mice to the optogenetic apparatus. Mice that failed to reach criteria within four days were given an additional 1-2 days of CRF training. Subsequent behavioral trials began with acute anesthetization with isoflourane and connection to fiber optic leads prior to training. Following CRF training, mice experienced three days of a variable-interval (VI) 30 task, in which they were rewarded on average 30 seconds (15-45 second range) contingent on lever pressing, followed by 8 days of VI60 training (rewarded every 60 seconds on average, ranging from 30 to 90 seconds, with each possible interval separated by 6 sec) (Nadel et al., 2020). VI sessions ended after 60 min or when 50 rewards had been earned. To assess the contribution of patches to habit formation, mice received optogenetic stimulation (5 mW, 5 Hz, 190 ms pulse width, 3 sec duration, see below) of patch neurons or terminals during the first headentry following each reward delivery in all VI60 trials. Patch activity is linked to reward-predicting cues or during reward consumption (Bloem et al., 2017; Yoshizawa et al., 2018), thus this stimulation timing was selected to modulate ongoing activity in patch neurons.

### Fiber Optic Implants

Fiber optic implants were custom fabricated and were comprised of 0.39 NA, 200 μm core Multimode Optical Fiber (ThorLabs) inserted into a multimode ceramic zirconia ferrules (1.25mm OD, 230um ID; SENKO). The fiber optic was affixed in the ferrule with two-part epoxy (353ND; Precision Fiber Products). Each end of the fiber optic was polished using fiber optic sandpaper (ThorLabs) and functionality was tested ensuring minimal loss of light power and even output prior to implantation.

### Laser Stimulation

Mice received blue laser stimulation (473 nm, 5 mW, 5 Hz, 190 ms pulse width, 3 sec duration) from a diode-pumped single-state laser (Laserglow) which was connected via fiber optic (Doric Lenses) to a commutator (1×2 Fiber-optic Rotary Joint) allowing for free rotation and splitting of the beam (Doric Lenses). The commutator was connected to two fiber optic leads, which were attached bilaterally to ferrules on fiber optic implants with a ceramic sleeve (Precision Fiber Products). Laser output was calibrated to 5 mW from the end of fiber optic leads before training each day using an optical power meter (ThorLabs). Laser parameters were the same for all behavioral tasks (VI60, Laser Probes, Open Field) with the exception of Real-Time Place Preference, where laser stimulation duration was cycled 5 sec ON, then 5 sec OFF (see below).

### Probe Tests

Following 8 days of VI60 training, a reward devaluation test was conducted. Here, mice were given free access to sucrose for one hour prior to testing. Mice were individually caged during this access and all mice were observed to ensure they consumed sucrose. To quantify sucrose consumption, a subgroup of mice had sucrose bottles weighed before and after free access. After the pre-feeding session, mice were given a 5-min probe test in which the lever was extended and presses were recorded, but no rewards were delivered. Reward devaluation is commonly used to probe habitual responding, and mice that persist in lever pressing during devaluation probes are considered more habitual (Adams and Dickinson, 1981; Gremel and Costa, 2013; O’Hare et al., 2016). Following devaluation probes, mice experienced one day of VI60 training to reinstate habitual responding. The following two days, mice were also tested on a 60 minute omission probe test in which the action-outcome contingency was reversed. Here, mice had to refrain from pressing the lever for 20 seconds to obtain a reward, and pressing the lever reset the timer (Nadel et al., 2020). This probe was employed as a second metric of habitual responding, as habitually responding mice are slower to reverse learned action-outcome contingencies (Yu et al., 2009). Following two days of omission trials, mice were again retrained on a VI60 schedule to reinstate lever pressing. The following day, mice underwent a “laser probe” trial. Here, mice again underwent reward devaluation by gaining free access to sucrose for one hour (as described above).

Mice were then returned to operant chambers and the lever was extended and presses were recorded, but no rewards were delivered. At variable intervals between 30-90 sec (6 sec between each possible interval) laser stimulation was delivered to fiber optic implants (5 mW, 5 Hz, 190 ms pulse width, 3 sec duration), and laser probe trials lasted a total of 30 min. This probe was conducted to determine the acute effects of patch stimulation on responding following reward devaluation.

### Real-Time Place Preference

Following operant conditioning tasks, mice were returned to *ad libitum* access to homecage chow. At least 3 days later, fiber optic implants were again connected to fiber optic leads and mice were placed in a 2-chamber place preference apparatus (Med Associates). Each chamber was 16.8 cm L x 12.7 cm W x 12.7 H with opaque walls. Chambers were distinguishable based on different flooring (grid vs bars) and different wall coloring (white vs black), and the orientation of the chamber did not change across place preference trials. To allow fiber optic movement and prevent mice from exiting the chamber, a custom, clear plexiglass wall extension (45.7 cm tall, 58.4 cm total height) was placed on the walls above the behavioral apparatus and no lid was utilized. Mice underwent two days of real-time place preference trials. Here, one chamber was randomly selected to trigger laser stimulation when mice entered or remained in the ‘active’ chamber, and the active chamber was counterbalanced across days. Location in the chambers was monitored by 12 evenly-spaced infrared beam breaks located near the floor of the apparatus. At the first occurrence of a beam break on the active side, laser stimulation was delivered to the fiber optic implants (5 mW, 5 Hz, 190 ms pulse width). As striatal stimulation can result in freezing depending on the neuronal population activated (Kravitz et al., 2010), laser stimulation was cycled ON for 5 sec and OFF for 5 sec. This pattern of stimulation occurred until a beam break occurred in the inactive chamber, when stimulation was halted until the next beam break in the active chamber. Time spent on either side was compared and averaged across each day to account for inherent preferences for either side. This task was performed to determine if optogenetic patch stimulation was inherently reinforcing, as suggested by a previous electrical self-stimulation experiment (White and Hiroi, 1998).

### Open Field

At least one day following RTCPP trials, fiber optic implants were again connected to fiber optic leads and mice were placed in an open field apparatus (42 cm wide x 42 cm long x 30 cm tall) to determine the effects of acute patch stimulation on locomotor activity. Every 60 sec laser stimulation (5 mW, 5 Hz, 190 ms pulse width, 3 sec duration) was delivered to implants. Mouse locomotion was monitored by a camera and analyzed online using Bonsai software (Open-Ephys). Movement was detected using a contrast-based binary region analysis and extraction of location in the video frame (Lopes 2015 Frontiers in Bioinformatics).

### Fast-Scan Cyclic Voltammetry

To determine the impact of patch activation on striatal dopamine release, we utilized fast-scan cyclic voltammetry (FSCV) to monitor real-time changes in striatal dopamine levels while simultaneously activating patch terminals with optogenetics *in vivo*. Fast-scan cyclic voltammetry was performed using custom glass-sealed, carbon-fiber microelectrodes (Cahill et al., 1996; Howard et al., 2011). Recordings were made by applying a triangular waveform (0.4 to 1.3 V and back, 400 V/s) every 100ms to the exposed tips of carbon-fiber microelectrodes. Voltammetry and stimulus control was performed by a WaveNeuro potentiostat (Pine Research) and was computer-controlled using HDCV software, which was generously provided by the Chemistry Department at UNC (Bucher et al., 2013). A subset of Sepw1-NP67 mice were injected with AAV driving Cre-dependent expression of ChR2 as described above. At least 3 weeks later, these mice were anesthetized using urethane (1 g/kg i.p. delivered in 2 injections separated by ~20 min) and placed in a stereotactic apparatus. An incision was made in the scalp and holes were drilled above the dorsal striatum (+0.8 AP, ±1.5 ML), SNc (−3.2 AP, ±1.5 ML), and pedunculopontine tegmental nucleus (PPTg, −0.68 AP from lambda, ±0.7 ML). The PPTg sends excitatory projections to dopamine neurons and was targeted with electrical stimulation to elicit dopamine release in the striatum (Forster and Blaha, 2003; Zweifel et al., 2009). An Ag/AgCl reference electrode was affixed in the superficial cortex. A carbon-fiber microelectrode was placed in the dorsal striatum (−2.3 DV), and during implantation the carbon-fiber was cycled at 60 Hz to allow the electrode to equilibrate and switched to 10 Hz ~20 min prior to data acquisition. A twisted bipolar stimulating electrode (Plastics One, Roanoke, VA, USA) connected to a DS4 Biphasic Constant Current Stimulus Isolator (Digitimer) was lowered in 0.1-mm increments starting at −1.5 DV into PPTg until robust dopamine increases were detected in the dorsal striatum. Stimulus trains consisted of 60 biphasic pulses delivered at 60 Hz at a current of 400-600 μA and was synchronized with recordings so that sampling and stimulation did not overlap. Stimulation intensity varied across subjects to elicit robust dopamine release but was fixed at the beginning of data collection and did not alter thereafter. Once stable dopamine release was detected, a fiber optic cable was inserted above SNc (−2.0 DV) to target patch terminals and was incrementally lowered to optimize placement (see Figure 6A for graphic of experimental design). Optogenetic stimulation consisted of 1 sec pulses of blue laser light delivered at 5-10 mW. Three trial types were then conducted: 1. Electrical stimulation of PPTg alone (“E stim” trials), 2. Optogenetic stimulation of patch terminals in SNc (“Opto” trials), or 3. Simultaneous electrical stimulation of pedunculopontine tegmental nucleus and optogenetic activation of patch terminals (“Both” trials). The order of trials was selected randomly until one of each trial type had been collected, then this process was repeated a minimum of 3 times. All recordings were separated by at least 3 minutes to avoid neurotransmitter vesicle depletion.

### Histology and Microscopy

At the cessation of all behavioral tests, mice were deeply anesthetized with isoflurane (4%, 2 l/min O2) and transcardially perfused with 0.9% saline and 4% paraformaldehyde (PFA). Brains were removed and allowed to post-fix in 4% PFA at 4°C for at least 24 h. Brains were then transferred to a 30% sucrose solution and returned to 4°C for at least 48 h. Brains were sectioned on a freezing microtome into 20 μm sections. A subset of striatal sections from optogenetic experiments were mounted and imaged to determine ChR2 expression. A separate set of sections from Sepw1-NP67 mice were washed 3X in Tris buffered saline (TBS) and blocked in 3% horse serum and 0.25% Triton X-100 prior to antibody staining. Sections were then incubated in a 1:500 dilution of anti-μ-opioid receptor polyclonal rabbit antibody (Immunostar, cat #24216) for 24–48 h at 4°C. A separate set of tissue was procured from Sepw1-NP67 mice crossed to a Cre-dependent GFP-reporter line to characterize Cre expression. This tissue was processed as described above, but was incubated in a 1:500 dilution of anti-μ-opioid receptor polyclonal rabbit antibody (Immunostar, cat #24216) and anti-GFP polyclonal guinea pig antibody (Synaptic Systems, cat#132–004) for 24–48 h at 4°C. Tissue was visualized using a Leica DM4000B fluorescent microscope or a Zeiss LSM 880 confocal microscope.

### Data Analysis

Mean and normalized press and head entry rates were compared across training and probe trials. As press rates in mice with lesioned patches have been shown to be variable across training days (Nadel), press and entry rates were normalized to average response rate across all VI60 trials to compare to probe trials. Omission and laser probe press and entry rates were normalized to the reinstatement VI60 training before each probe trial. We expected potentially opposing effects of modulating differing terminal sites, but across VI60 training, devaluation probe trials, omission, laser probe trials, open field, and place preference tasks we noted no statistical differences between different fiber optic implantation sites in ChR2 groups (Supporting Figures 1–5), therefore, groups were collapsed and comparisons were made between ChR2 mice and YFP controls. The ratio of time spent in active:inactive chambers was averaged across two days of the place preference task and then averaged across groups. Velocity in the open field was calibrated from megapixels/frame to cm/sec using Matlab software MATLAB (R2018b, Mathworks). Press and entry rates were calculated using Excel (Microsoft). Autocorrelations, cross-correlations, and real-time press and entry rates in laser probes, were determined using custom scripts written in MATLAB (R2018b, Mathworks). To control for individual differences in baseline responding and to determine laser-induced changes in responding, laser probe press and entry rates were subtracted from baseline responding 10 sec before laser onset before being averaged. To quantify responses in laser probe trials, response rate was averaged across 1 sec just prior to and 5-8.5 sec following laser onset. FSCV data was analyzed in HDCV (UNC Chemistry Department). Voltammetric current vs time and current vs. voltage traces were collected and averaged for each trial type within experiments (see above) before being averaged between subjects. Evoked amplitudes were normalized to maximum current in PPTg stimulation only trials (‘E stim’ trials) to account for different amplitudes of dopamine responses across subjects.

### Statistical Analysis

Statistical analysis was performed by GraphPad Prism 7.04 (GraphPad) or Matlab (R2018b, Mathworks). Press and entry rates during VI60 and omission probes were compared using Two-Way Repeated Measures ANOVA with *post hoc* Sidak multiple comparison tests. Comparisons between stimulation sites, evoked dopamine responses, and sucrose consumption between groups were compared using One-Way ANOVA with *post hoc* Tukey’s or Holm-Sidak multiple comparison’s tests. Press rates in VI60, VI30, and devaluation probes, as well as time on stimulation side in place preference, baseline velocity, changes in velocity, and autocorrelations were compared using unpaired student’s t-tests. Changes in press and entry rates in laser probe trials were compared using paired student’s t-tests. Pearson’s Correlation was utilized for all correlations. Statistical outliers were determined using the ROUT (robust regression followed by outlier identification) method (Q=0.5%) in GraphPad Prism 7.04 (GraphPad) and were removed prior to statistical analyses. Finally, mice lacking ostensible viral expression in the striatum were excluded prior to analysis. For all tests significance was defined as p ≤ 0.05.

## Acknowledgements

The authors would like to thank Drs. Charles Gerfen (National Institute of Mental Health) and Nathaniel Heintz (The Rockefeller University) for generously providing Sepw1 NP67 mice. This work was supported by NIH grant 1R15MH122729-01. J.A.N. was supported by the Nu Rho Psi Undergraduate Research Grant and the Robert Rich Student Research Grant through Oberlin College. Finally, the authors would also like to thank Lori Lindsay, Forrest Rose, Dorothy Auble, Gigi Knight, Bill Mohler, Chris Mohler and Laurie Holcomb for research support.

## Supporting Figures

**Supporting Figure 1.**
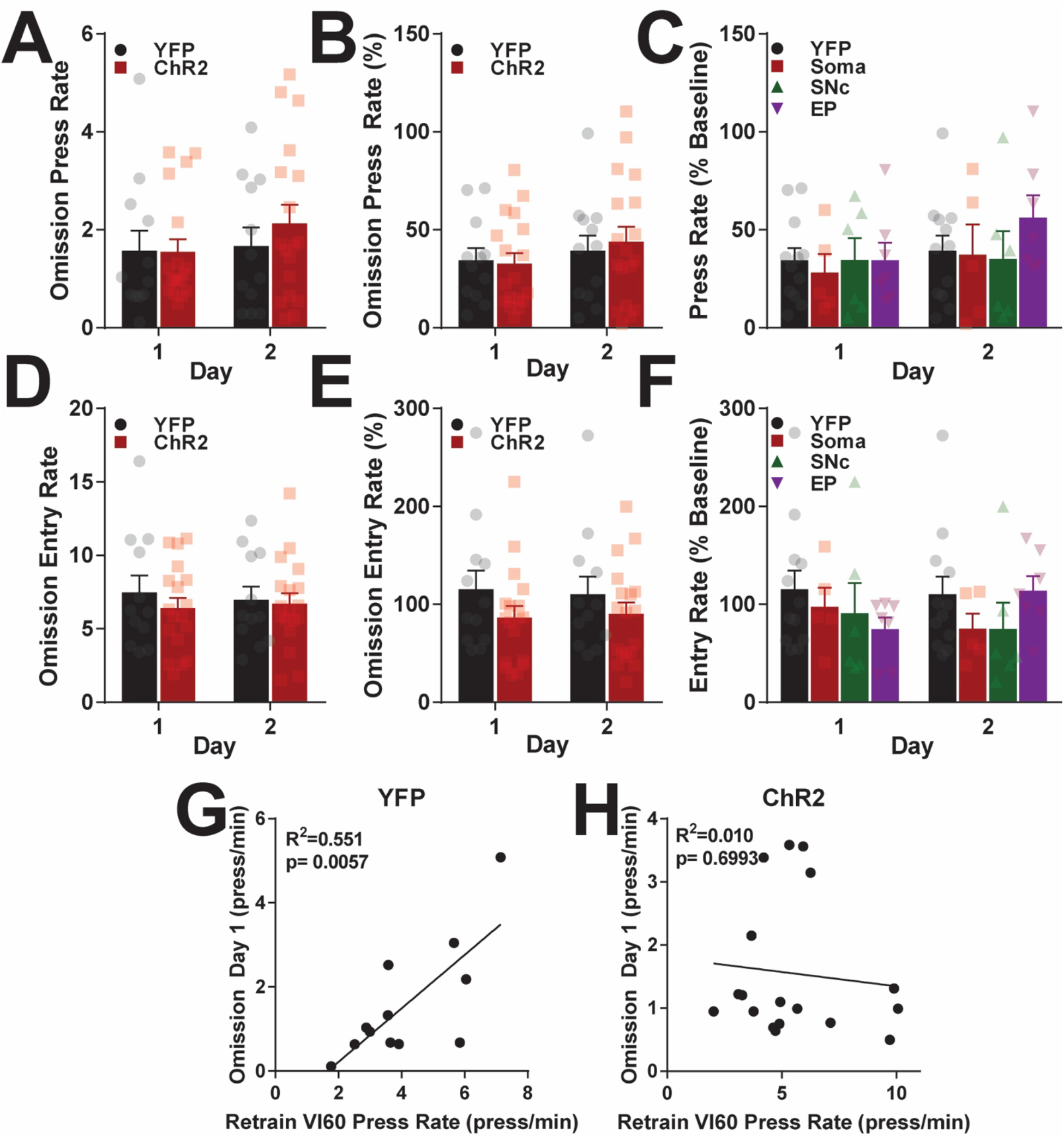
Effects of optogenetic patch stimulation during learning on omission. A. Average press rates during omission probes for YFP control and ChR2 mice across days (two-way repeated measures ANOVA, no significant effects of time, group, or interaction). B. Average press rates normalized to responding during VI60 retraining during omission probes (two-way repeated measures ANOVA, no significant effects of time, group, or interaction). C. Same data as B, but broken into fiber optic placement groups (two-way repeated measures ANOVA, significant effect of time, F_(1,26)_ = 4.56, p = 0.042), no significant effect of group or interaction). D. Average head entry rates during omission across days (two-way repeated measures ANOVA, no significant effects of time, group, or interaction). E. Average entry rates normalized to baseline entry rates in VI60 retraining (two-way repeated measures ANOVA, no significant effects of time, group, or interaction). F. Same as E, but broken into fiber optic placement groups (two-way repeated measures ANOVA, significant time x group interaction, F_(3,26)_ = 3.87, p = 0.021, no significant *post hoc* Tukey tests). G-H. Correlation of omission press rate on day 1 vs. VI60 retraining day immediately preceding omission for YFP (G) or ChR2 (H) mice. Data are mean ± SEM.

**Supporting Figure 2.**
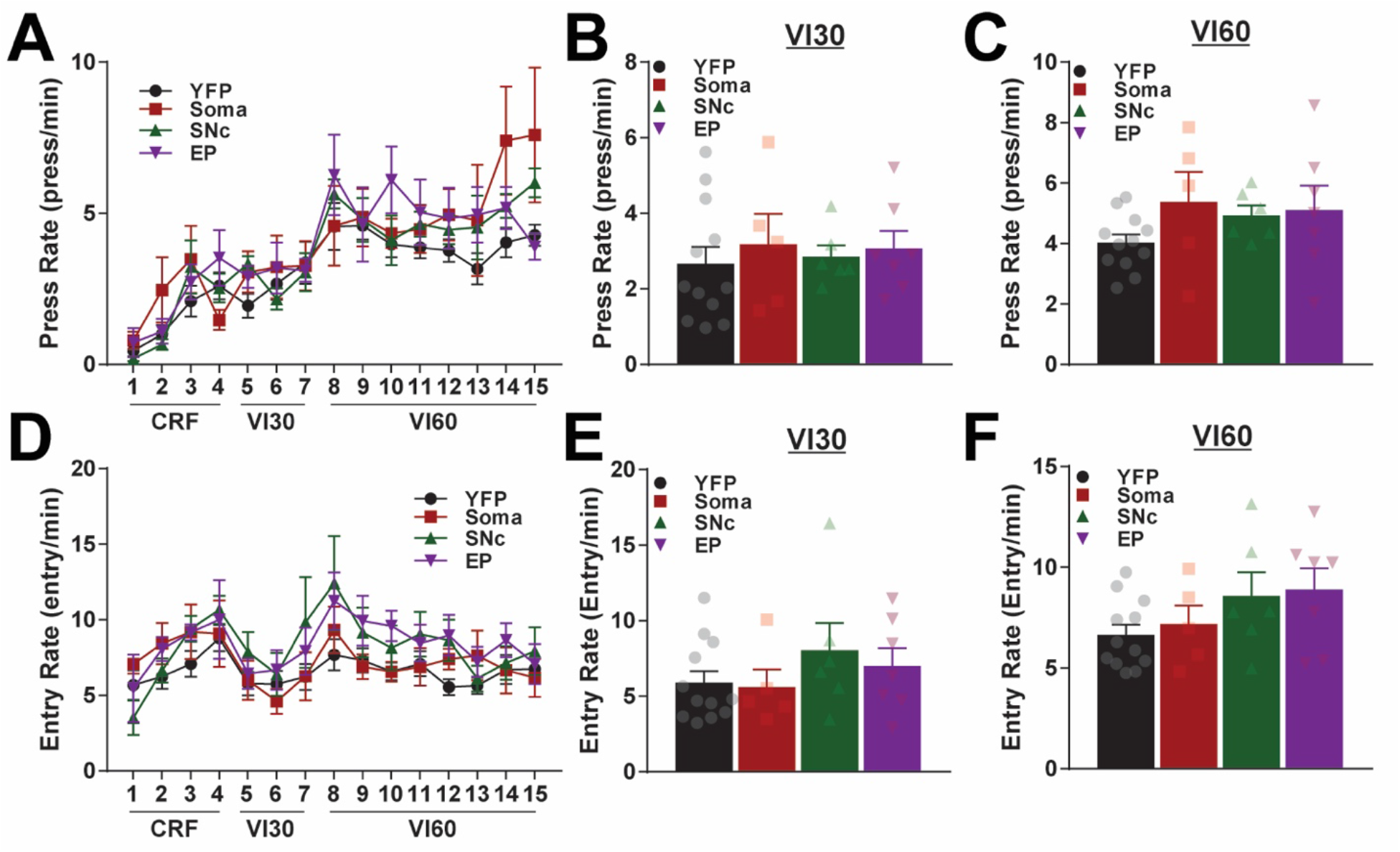
Optogenetic patch stimulation during variable interval training by implantation site group. A. Average press rates across continuous reinforcement (CRF), variable interval 30 (VI30), and variable interval 60 (VI60) training by day (two-way repeated measures ANOVA, significant effect of time, F_(14,364)_ = 21.98, p < 0.0001, no significant effects of group or interaction). B-C. Average press rates across all VI30 (B; one-way ANOVA, F_(3,26)_ = 0.21, p = 0.89) and VI60 (C; one-way ANOVA, F_(3,26)_ = 1.34, p = 0.28) days. D. Average head entry rates across training by day (two-way repeated measures ANOVA, significant effect of time, F_(14,364)_ = 6.12, p < 0.0001, no significant effects of group or interaction). E-F. Average entry rate across all VI30 (E; one-way ANOVA, F_(3,26)_ = 1.78, p = 0.18) and VI60 (F; one-way ANOVA, F_(3,26)_ = 0.79, p = 0.51) days.

**Supporting Figure 3.**
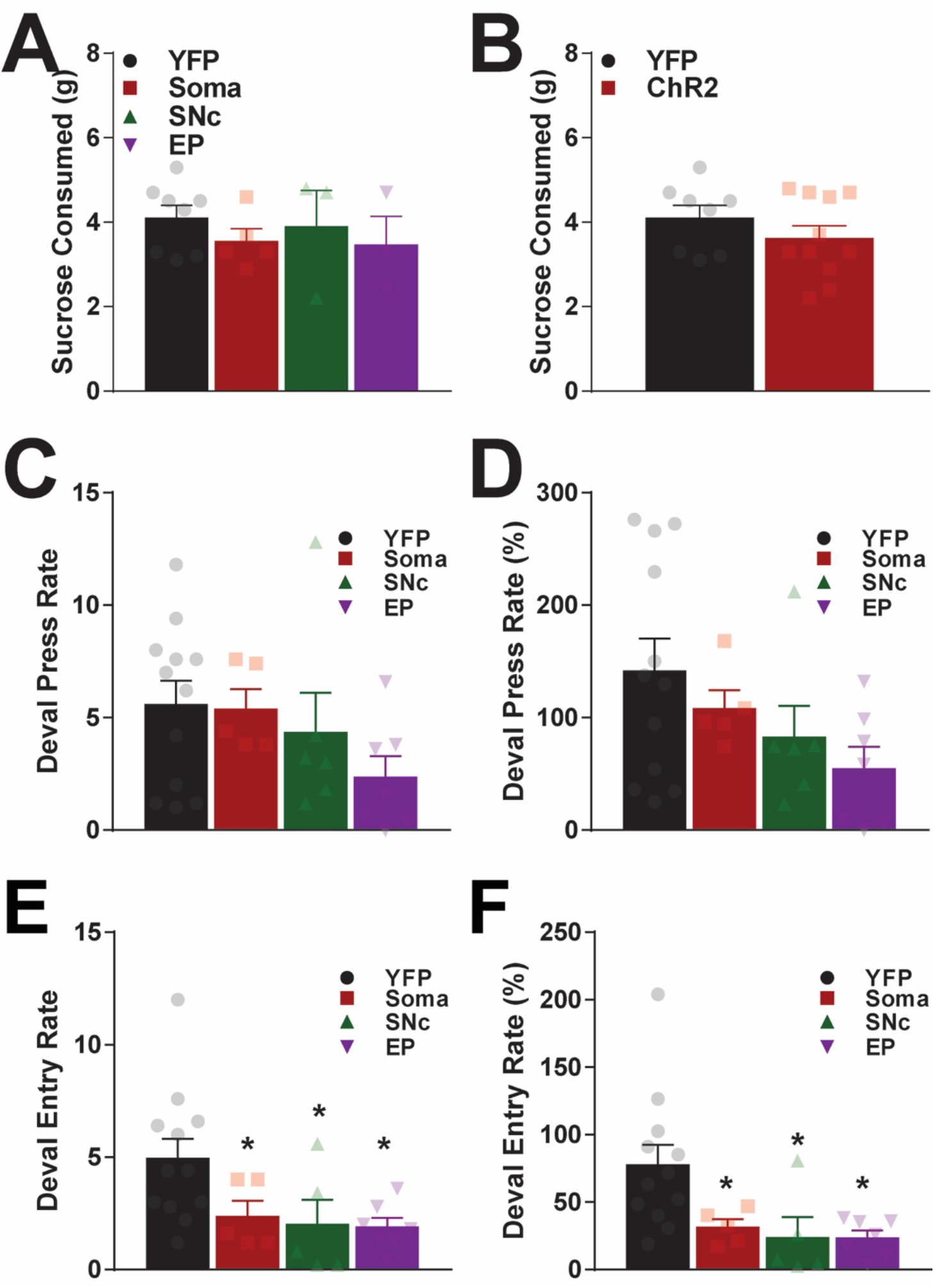
Sucrose consumed during free access and press and entry rates in devaluation by implantation site group. A-B. Average sucrose consumed during 1h free access by implantation site group (A; oneway ANOVA, F_(3,15)_ = 0.531, p = 0.67) and collapsed into YFP/ChR2 groups (B; unpaired t-test, t_17_ = 1.17, p = 0.26). C-D. Average press rate (C; one-way ANOVA, F_(3,26)_ = 1.53, p = 0.23) and average press rates normalized to responding across VI60 training (D; one-way ANOVA, F_(3,26)_ = 2.17, p = 0.12). E-F. Average head entry rates (E; one-way ANOVA, F_(3,25)_ = 3.626, p = 0.027) and entry rates normalized to baseline VI60 responding (F; one-way ANOVA, F_(3,25)_ = 4.644, p = 0.010). Data are mean ± SEM. *significant Holm-Sidak *post hoc* test

**Supporting Figure 4.**
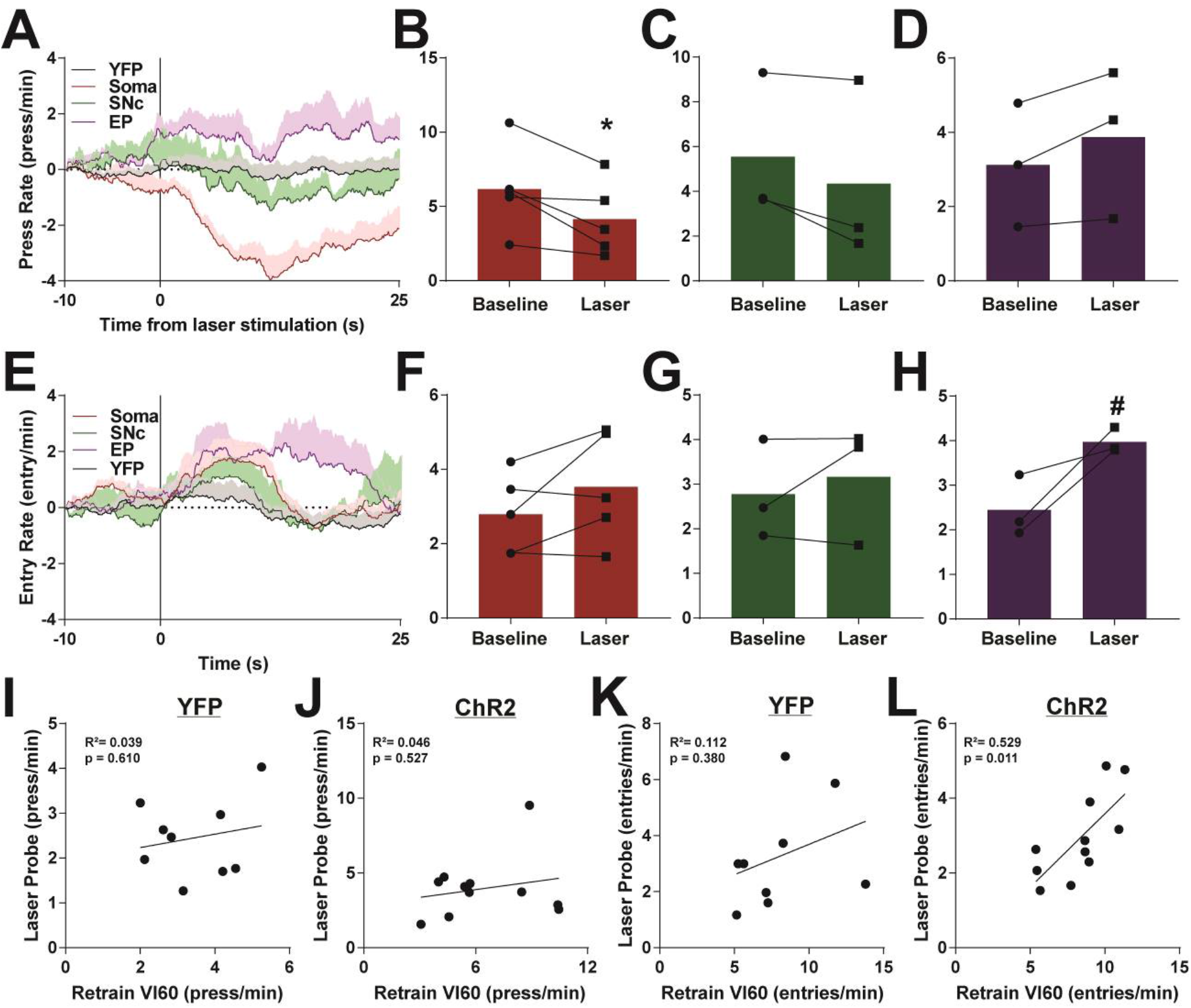
Effects of acute laser stimulation during laser probe trials by implantation site group. A. Average lever press rate before and after laser onset (vertical line; 3 sec, 5 Hz stimulation). B-D. Average press rates before and following laser onset for Soma (B; paired t-test, t_4_ = 3.10, p = 0.036), SNc (C; paired t-test, t_2_ = 2.48, p = 0.13), and EP (D; paired t-test, t_2_ = 2.58, p = 0.12) groups. E. Average head entry rates before and after laser onset. F-H. Average entry rates before and after laser onset for Soma (F; paired t-test, t_4_ = 1.69, p = 0.17), SNc (G; paired t-test, t_2_ = 0.78, p = 0.51), and EP (H; paired t-test, t_2_ = 3.24, p = 0.084) groups. I-J. Correlation of lever press rates during laser probe trials and VI60 retraining the day before for YFP (I) and ChR2 (J) mice. K-L. Correlation of head entry rates during laser probe and VI60 retraining for YFP (K) and ChR2 (L) mice. Data are mean ± SEM. *#P* < 0.1

**Supporting Figure 5.**
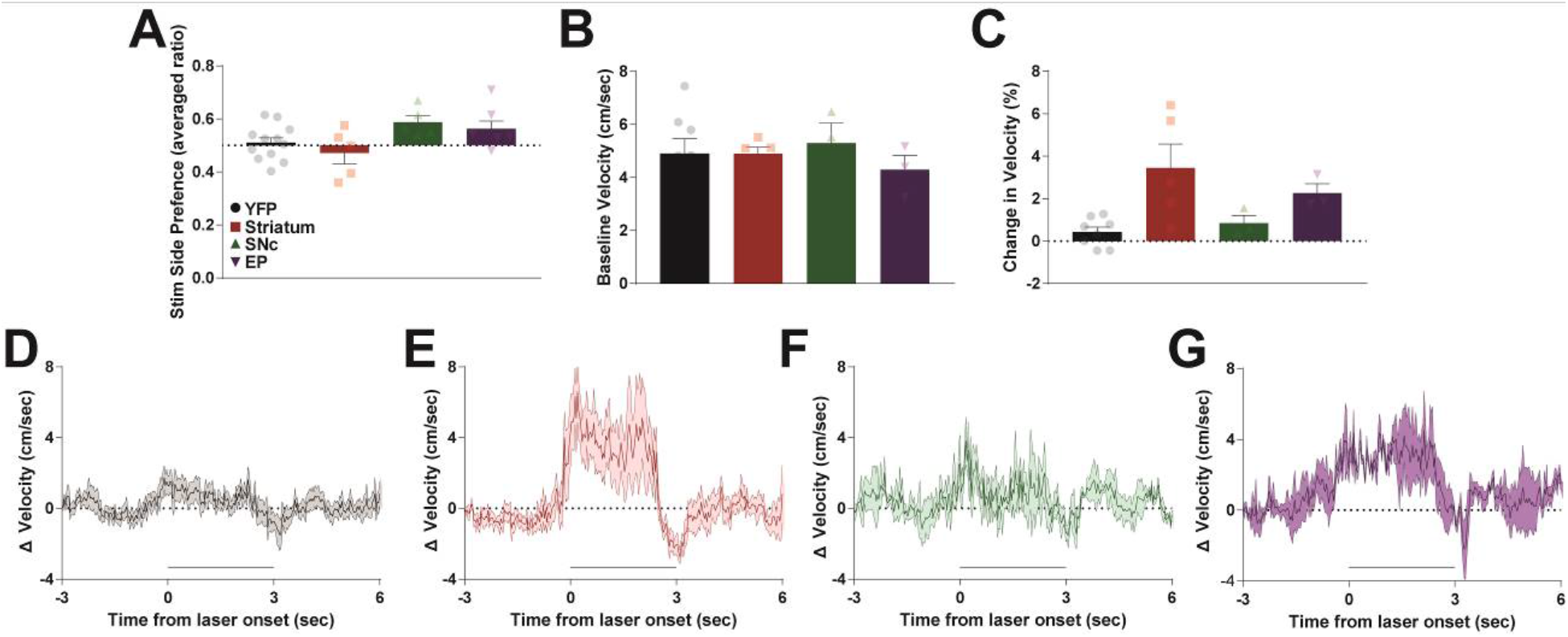
Effects of optogenetic stimulation of patches on reinforcement and locomotion by implantation site groups. A. Time spent on stimulated side of a two-chamber place preference apparatus. Stimulation side was counterbalanced across two days and averaged between days (one-way ANOVA, F_(3,25)_ = 3.023, p = 0.048, no significant *post hoc* Holm-Sidak tests). B. Baseline velocity in open field (one-way ANOVA, F_(3,15)_ = 0.33, p = 0.80). C. Change in velocity following laser onset in open field (one-way ANOVA, F_(3,15)_ = 5.22, p = 0.012, no significant *post hoc* Holm-Sidak tests). D-E. Average baseline normalized velocity before and after laser onset (5 Hz, 3 sec; denoted by thick black line) for YFP (D), Soma (E), SNc (F), and EP (G) mice. Data are mean ± SEM.

**Supporting Figure 6.**
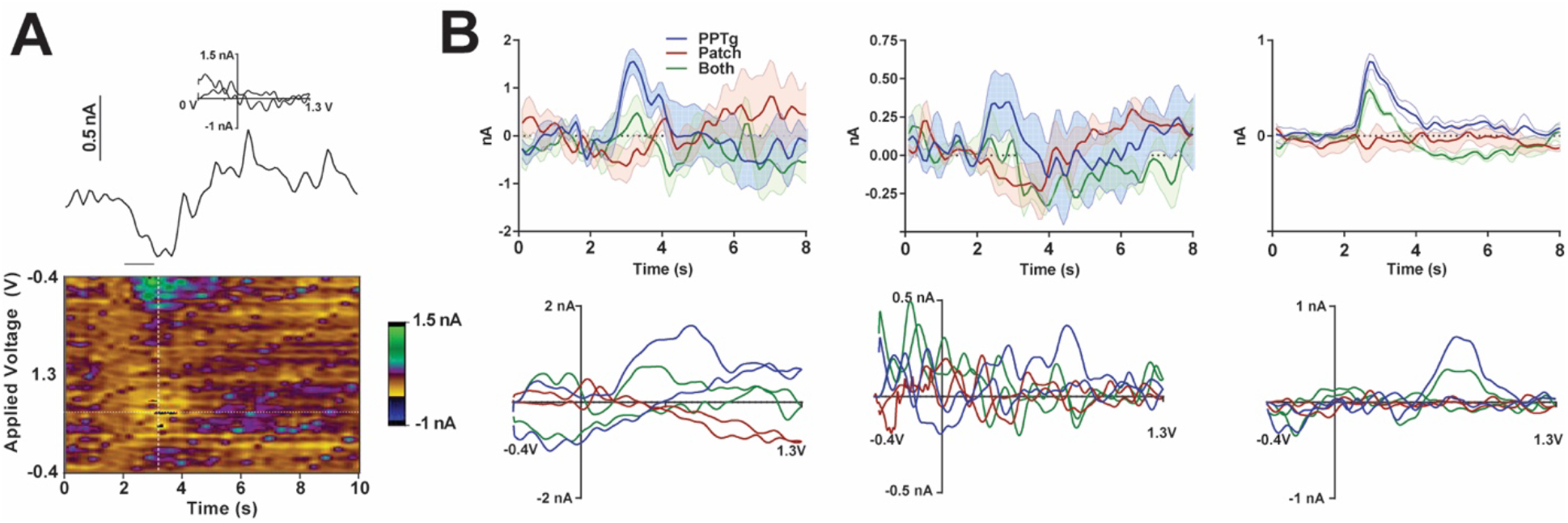
Average data from each replicate in FSCV experiments. A. Representative decrease in current recorded during an “opto” trial (stimulation of patch terminals only). The line shows recorded current relative to stimulation delivery (straight line below current trace) above a pseudo-color plot. The color plot shows current collected (in color) at each waveform scan (y-axis) and across time (x-axis). INSET: a “cyclic voltammogram” collected at the vertical white dotted line on the pseudo-color plot. B. Average current (top) and cyclic voltammograms (bottom) recorded across three trials for each of the three stimulation conditions. Each current and cyclic voltammogram plot is from an individual FSCV experiment.

